# Fronto-medial theta coordinates posterior maintenance of working memory content

**DOI:** 10.1101/2021.03.18.435966

**Authors:** Oliver Ratcliffe, Kimron Shapiro, Bernhard P. Staresina

## Abstract

How does the human brain manage multiple bits of information to guide goal-directed behaviour? Successful working memory (WM) functioning has consistently been linked to oscillatory power in the theta frequency band (4-8 Hz) over fronto-medial cortex (fronto-medial theta, FMT). Specifically, FMT is thought to reflect the mechanism of an executive sub-system that coordinates maintenance of memory contents in posterior regions. However, direct evidence for the role of FMT in controlling specific WM content is lacking. Here we collected high-density Electroencephalography (EEG) data whilst participants engaged in load-varying WM tasks and then used multivariate decoding methods to examine WM content during the maintenance period. Higher WM load elicited a focal increase in FMT. Importantly, decoding of WM content was driven by posterior/parietal sites, which in turn showed load-induced functional theta coupling with fronto-medial cortex. Finally, we observed a significant slowing of FMT frequency with increasing WM load, consistent with the hypothesised broadening of a theta ‘duty cycle’ to accommodate additional WM items. Together these findings demonstrate that frontal theta orchestrates posterior maintenance of WM content. Moreover, the observed frequency slowing elucidates the function of FMT oscillations by specifically supporting phase-coding accounts of WM.

**Significance Statement:** How does the brain juggle the maintenance of multiple items in working memory (WM)? Here we show that increased WM demands increase theta power (4-8 Hz) in fronto-medial cortex. Interestingly, using a machine learning approach, we found that the content held in WM could be read out not from frontal, but from posterior areas. These areas were in turn functionally coupled with fronto-medial cortex, consistent with the idea that frontal cortex orchestrates WM representations in posterior regions. Finally, we observed that holding an additional item in WM leads to significant slowing of the frontal theta rhythm, supporting computational models that postulate longer ‘duty cycles’ to accommodate additional WM demands.

## Introduction

Working memory (WM) is the ability to retain and manipulate information over short delays (Baddeley, 1992). It is thought to rely on at least two functionally distinct sub-systems (Baddeley, 2003). The first is an executive system, which directs cognitive resources and oversees the prioritisation and readout of representations. This system interacts with the representational system, which directly holds task-relevant informational content. These two systems rely on disparate brain regions, in particular frontal and parietal areas (Bressler & Menon, 2010; Imaruoka et al., 2005; Owen et al., 2005; Vogel & Machizawa, 2004). Importantly, WM requires functional interactions between these systems. A prime mechanism to facilitate inter-regional communication in service of WM is oscillatory activity in the theta range (4-8 Hz) (Sauseng et al., 2010; Von Stein & Sarnthein, 2000).

Specifically, according to a framework put forward by Lisman & Jensen (2013), theta oscillations provide gated processing windows in which neural representations of target information (‘WM items’) become active. Consistent with this model, non-human primate work has demonstrated that neurons coding for item-related information preferentially fire during particular theta phases (Siegel et al., 2009). Similarly, a recent study using intracranial recordings in humans has shown nesting of stimulus-related gamma activity (>30 Hz) in specific theta phases (Bahramisharif et al., 2018). Theta oscillations may thus constitute the mechanism governing goal-directed interactions between frontal executive and parietal representational subsystems (D’Esposito & Postle, 2015).

WM tasks consistently induce theta power increases at fronto-medial sites (fronto-medial theta, FMT) (Hsieh & Ranganath, 2014; Michels et al., 2010; Raghavachari et al., 2001), originating from the medial prefrontal and anterior cingulate cortices (Meltzer et al., 2008; Tsujimoto et al., 2006). FMT has been linked to behavioural WM performance (Maurer et al., 2014), as well as to individual WM capacity (Zakrzewska & Brzezicka, 2014). Consistent with this role for FMT, transcranial magnetic stimulation (TMS) in the theta range improved WM performance (Riddle et al., 2020). Importantly, non-invasive electrophysiological studies have detected fronto-parietal coupling via theta oscillations during WM tasks (Payne & Kounios, 2009). Additionally, experimental induction of fronto-parietal theta coupling via transcranial alternating current stimulation (tACS) increased WM performance (Polanía et al., 2012; Reinhart & Nguyen, 2019). Finally, EEG latency analyses between frontal and parietal regions suggest that the frontal cortex is the driver of these inter-regional interactions (Sauseng et al., 2004).

These findings suggest the intriguing scenario that WM content, represented and maintained in posterior/parietal cortex, is orchestrated by frontal control mechanisms via theta oscillations. However, at present it is unclear (i) whether parietal cortex represents individual items held in WM and (ii) whether those parietal WM memoranda are in turn coordinated by FMT. Finally, it is unclear how theta rhythms orchestrate different WM loads, e.g., maintaining two instead of one item in WM. On the one hand, an increase in load may be accommodated by an increase in theta amplitude/power, reflecting the participation of larger neuronal assemblies (Buzsáki & Draguhn, 2004). Conversely, the theta-gamma framework referred to above emphasises a role of oscillatory phase, which allows for separation of individual chunks of information (Lisman & Jensen, 2013). Accordingly, increasing load might induce a slowing of an individual’s theta frequency - the slower the frequency, the more item-coding assemblies can fire within a given cycle.

To answer these questions, we designed a paradigm in which we systematically varied WM load and decoded the category of a maintained visual stimulus whilst recording high-density EEG. We first show that frontal theta power increases with WM load. Second, individual items held in WM were decodable from the EEG signal in parietal cortex. Intriguingly, the same channels that enabled decoding showed significant coherence in the theta band with the frontal channels identified previously. Finally, we found that theta frequencies slow with increased WM load, consistent with phase-coding accounts of WM maintenance. Together, these results elucidate the role of theta rhythms for linking frontal control mechanisms with parietal content representations during WM maintenance.

## Method

### Participants

Thirty-three participants in total were tested. Participants were right-handed, aged between 18 and 35, and had no history of psychological or neurological disorder. Data from two participants were removed for low behavioural performance (see Statistics section for further detail). Data from a further three participants were removed due to poor EEG quality. All analyses therefore focussed on the remaining 28 participants (18 female, mean age of 22.64 years, SD = 3.95, range = 18-33).

### Procedure

Two behavioural tasks were employed in this experiment: a delayed-match-to-sample (DMS) task and an n-back task featuring two levels of working memory load. Stimuli were 350×350 pixel colour images of one of three categories: object, face, or scene. There were five unique stimuli from each category. The DMS task included all three categories, whereas the n-back task used only the object and the scene stimuli. The additional category in the DMS task was included to facilitate alternative analyses outside the scope of the results we report here. The stimuli were obtained from the BOSS (Brodeur et al., 2010) and SUN (Xiao et al., 2010) online databases.

After EEG setup was complete, participants performed the first run of the DMS task. Within this run, each sample image (to-be-compared to the probe) was presented six times. For each trial a randomly selected probe image was presented. Following completion of the DMS task, participants completed the n-back task, which consisted of 12 blocks (8×2-back; 4×1-back). Each block contained 36+n trials. After completion of the n-back task, participants performed the second run of the DMS task in which each stimulus was again presented six times in a random order. The DMS task was run twice to maximise the number of trials per category. The two runs were performed before and after the n-back task to account for any changes in the EEG signal across the recording session (e.g., signal drift).

### Experimental Design and Statistical Analysis

The experimental tasks are illustrated in Figure 1. In the Delayed-match-to-sample (DMS) task, participants were asked to focus on a central fixation cross before an image of an object, scene, or face was presented for ≥750 ms. After a delay period of ≥2500 ms, a probe stimulus (randomly selected from the full stimulus set) was shown for ≥750 ms. In the subsequent response window, an ‘X’ was present on the screen for 750 ms and participants responded using either the left or right arrow key (counter-balanced across participants) to indicate whether the probe’s identity was the same as the first image presented in the trial (i.e., whether it was a ‘match’ or ‘non-match’). In both cases, participants were required to make a response. The duration of the initial stimulus, delay period, and the probe stimulus were all jittered so that trials lasted for the base duration plus 0, 50, 100, or 150 ms. Each trial’s temporal jitter was randomly assigned ensuring that each jitter possibility (including no jitter) was equally represented in each block independent of category.

**Figure 1.**
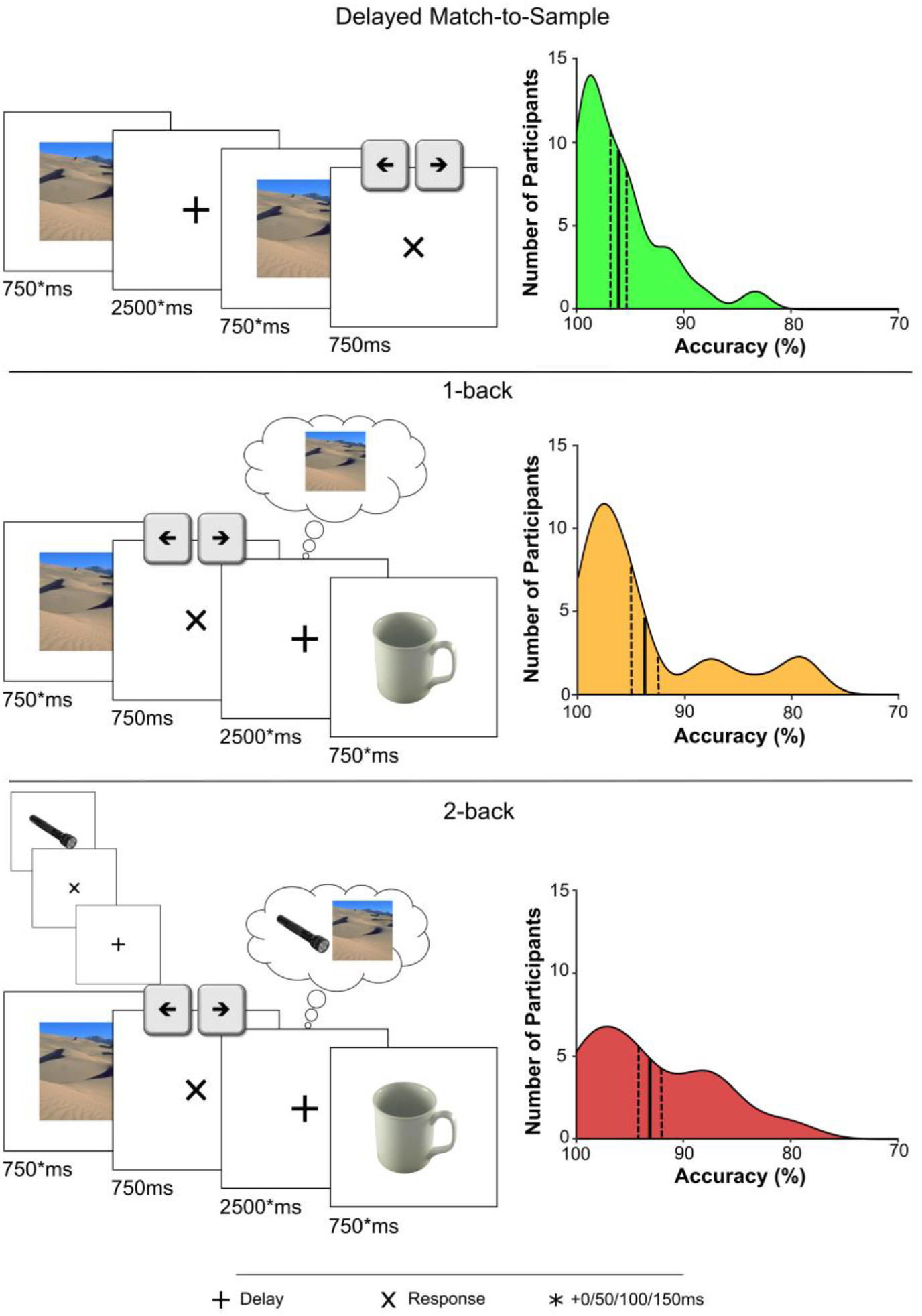
Experimental paradigm and behavioural results. Trial design (left) and distributions of behavioural accuracy (right) in three WM tasks performed during acquisition of EEG: a delayed-match-to-sample (DMS) task (top), a 1-back task (middle) and a 2-back task (bottom). Solid black lines indicate the mean behavioural accuracy and dashed black lines indicate the standard error of the mean across participants.

In the n-back task, participants were presented with an image of an object or a scene for ≥750 ms. An ‘X’ then appeared on the screen for 750 ms during which participants were required to respond ‘match’ or ‘non-match’ with either the left or right arrow key (counter-balanced across participants) to indicate whether the identity of stimulus just seen matched that of the stimulus seen *n* trials back. A ‘+’ was then presented for ≥2500 ms. Participants were required to maintain the relevant stimulus (1-back) or stimuli (2-back), so that they could make the n-back match/non-match judgement on the following trial. The stimulus and delay periods were jittered by 0, 50, 100, or 150 ms. As with the DMS task, the possible jitter options were balanced within blocks independent of stimulus category.

Behavioural accuracy was calculated as the proportion of correct responses out of all trials. For outlier analysis, a composite score for each participant was computed by taking the mean accuracy on all three tasks. Outliers were defined as any value more than 1.5 inter-quartile ranges below the lower quartile or above the upper quartile across all participants. As mentioned above, two participants were removed from subsequent analyses due to outlying low behavioural accuracy.

An alpha level of 0.05 was used as the threshold for statistical significance. Tests were conducted as 2-tailed except for testing classification accuracies against chance. In the latter case, as chance represents the floor of performance, a 1-tailed test was deemed more appropriate. To control for multiple comparisons across dimensions when assessing load-dependent power changes (channel/frequency/time) or when assessing searchlight classification (channel/time), non-parametric cluster-based permutation testing was employed (Maris & Oostenveld, 2007). This permutation testing was based on the maximum sum of a cluster’s t-values, 500 permutations and at least three neighbouring channels constituting a cluster.

### EEG setup and pre-processing

EEG data were collected using a BioSemi system with 128 channels at a 1024 Hz sampling rate. Data were re-referenced offline to the average of the two mastoids. Eye blinks were removed from data using independent components analysis, as implemented by ‘runica’ in Fieldtrip’s *ft_componentanalysis*. Consistently noisy channels were interpolated using a weighted average of neighbours. Data were then high-pass filtered at 0.3 Hz prior to all other analyses.

### Time-frequency calculations

Time-frequency spectra were calculated using Fieldtrip’s *mtmconvol* function. Power in frequencies from 2 to 10 Hz (0.5 Hz steps) were computed across the delay period using a Hanning taper. Power was resolved in 50 ms increments. Data were convolved with a variable number of cycles per frequency band. Two cycles were used for frequencies 2-3.5 Hz; 3 cycles for 4-4.5 Hz; 4 cycles for 5-5.5 Hz; and 5 cycles for 6-10 Hz.

Following this analysis, we assessed whether there was a difference in the FMT peak frequency for 1-back vs. 2-back tasks. To this end, spectral power was calculated for all channels that were members of the frontal cluster previously identified. Power was computed for frequencies between 4-8 Hz in 0.2 Hz increments across the full delay period for both conditions for each of these channels. For this analysis, power was calculated via Fieldtrip’s *mtmfft* function. Power spectra from individual channels were then averaged. Thus, for the resulting power spectrum of each participant and every trial, local maxima were identified (Matlab function *findpeaks*). The frequency at which the most prominent of these peaks occurred was logged for every trial. These peak frequency values were then separated into load conditions and averaged across trials. This value was obtained for each participant for 1-back and 2-back trials. Differences in theta peak frequency between the two load conditions were compared via a paired-samples t-test. To obviate the possibility that any difference in these values reflects a shift in the slope of the 1/f component of the EEG signal (Donoghue et al., 2020), the IRASA method (Irregular Resampling Auto-Spectral Analysis; (Wen & Liu, 2016)) was employed to remove the 1/f component from the signal. Additionally, to ensure that this result was not a consequence of any volatility in single-trial power spectra, the analysis was also conducted on smoothed frequency spectra. The spectra were smoothed prior to peak detection via a sliding mean average using the Matlab function *smoothdata*. The degree of smoothing was varied between 2 and 5 elements (approximately 0.2-1.0 Hz window).

### Classification

To decode object vs. scene representations during WM maintenance, multivariate pattern analyses (MVPA) was performed with the MVPA-light toolbox (Treder, 2020). To reduce computational time of classification, data were resampled to 200 Hz. Prior to classification, data were smoothed with a running average (100 ms sliding window) and baseline-corrected to the preceding 200 ms. For all classification analyses, linear discriminant analysis (LDA) was employed by taking the EEG channels as features at every time point. Classification was performed on correct trials only using a k-fold cross-validation procedure in which data were divided into 5 folds (4 training and 1 testing) and this was repeated 5 times. The accuracy values across folds and repetitions were then averaged to produce the final classifier performance.

To determine which channels were most informative to correct classification, we implemented a searchlight approach in which, moving around the channel map of all 128 channels successively, an individual channel and its neighbours (radius = 0.10) were used to classify the data. The resultant accuracy for each of the searchlight centres was then tested against chance. Searchlight classification was performed by training and testing during the window in which significant above-chance decoding was observed including all channels (880-1305 ms into the maintenance period, see below).

### Connectivity

To assess load-dependent changes in functional connectivity, cross-spectral densities were computed across the full time period via the Fieldtrip function *mtmconvol* using the same settings as in the time-frequency decomposition described previously. Pairwise channel coherence in the theta range (4-8 Hz) between channel Fz and every other channel was derived in the delay period of the 1-back task and the DMS task. Data from the DMS task were sub-sampled 10 times to accommodate for the lower trial count in the 1-back condition (Bastos & Schoffelen, 2016). These sub-sampled coherence maps were averaged before condition contrasts. Coherence values were compared statistically between 1-back and DMS conditions via cluster-based permutation tests in the time during the delay period where there was significant decoding above chance (880-1305 ms) (Maris & Oostenveld, 2007).

### Code

All behavioural tasks were created and presented using Matlab 2016b (MATLAB and Statistics Toolbox Release 2016b, The MathWorks, Inc., Natick, Massachusetts, United States) and Psychtoolbox (Version 3.0.16; Brainard, 1997; Kleiner et al., 2007). Analyses were conducted with custom Matlab and R scripts (Version 3.4.3; R Core Team, 2018). EEG analyses were performed with Fieldtrip functions (Version 20210308; Oostenveld et al., 2011). Plots also made use of *boundedline* (Kearney, 2019) and *hline and vline* (Kuczenski, 2019).

### Code Accessibility

The analysis scripts & processed data will be made available on the Open Science Framework (OSF). Raw data will be available upon reasonable request.

## Results

### Oscillatory Mechanisms of WM Maintenance

To test our main hypotheses regarding the role of fronto-medial theta (FMT) in WM, we focused on the comparison of the 1-back vs. the DMS task. The 2-back task was used to test the specific hypothesis on theta slowing as a function of WM load. First, we aimed to corroborate previous findings that engagement in WM-dependent tasks is supported by an increase in FMT power (Hsieh & Ranganath, 2014). Analysis of behavioural accuracy confirmed increased task difficulty for the 1-back task compared to the DMS task, despite high performance in both tasks (Figure 1; 1-back: 94% mean accuracy; SD = 6.59%; DMS: 96% mean accuracy; SD = 3.97%, paired-samples t-test, t_(27)_ = −2.40; p = 0.024). To examine oscillatory activity, spectral power was calculated in the delay period of both the 1-back and the DMS tasks, averaging power values across the full 2.5 second delay period. A permutation-based cluster-corrected paired-samples t-test was then conducted on these channel-frequency spectra. Results revealed a significant cluster in which theta power was greater in the 1-back task as compared to the DMS task. This cluster was centred primarily around 5.5-7 Hz but spanned the entire theta frequency band (4-8 Hz; Fig. 2A). Collapsing across the 4-8 Hz frequency range illustrates this effect is driven by fronto-medial channels (Fig. 2B), consistent with previous work identifying fronto-medial sources of WM-related theta oscillations (Onton et al., 2005; Zuure et al., 2020).

**Figure 2.**
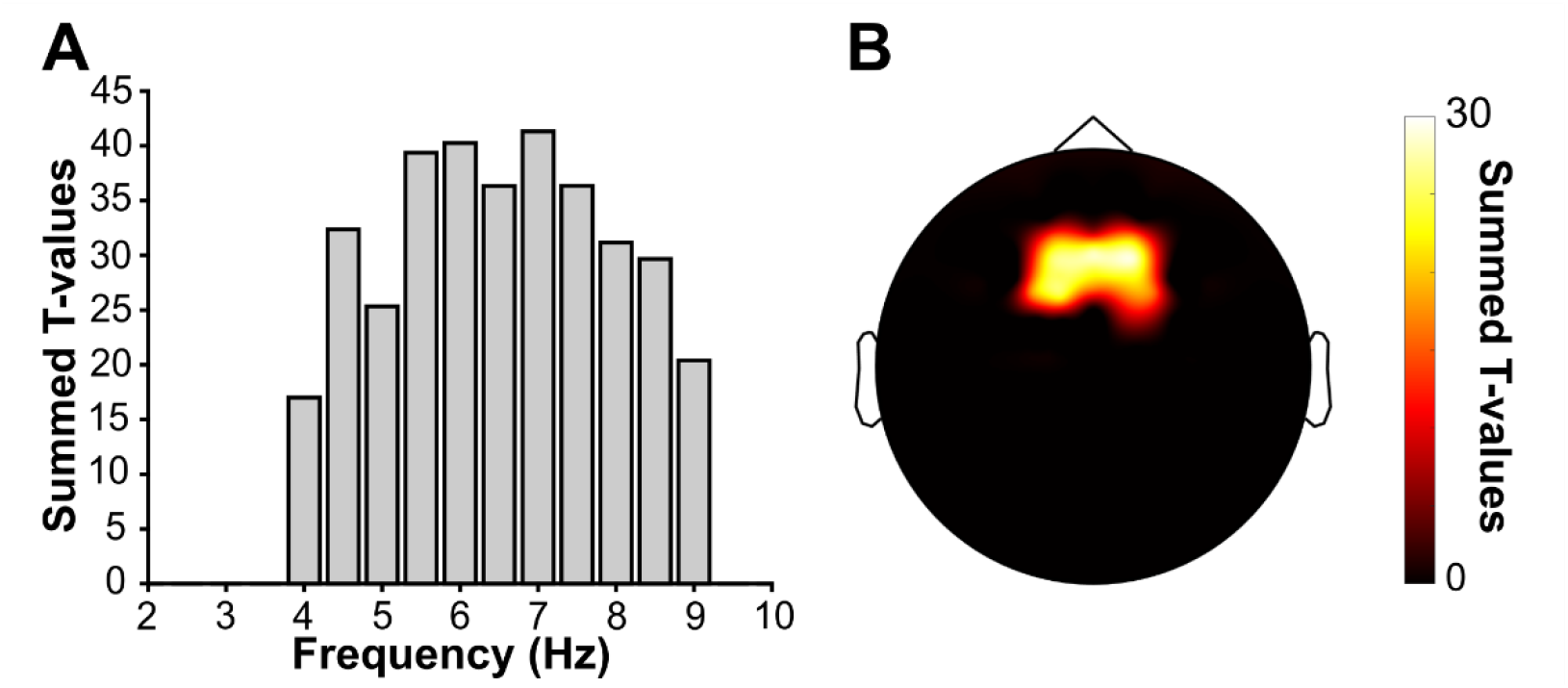
WM-induced fronto-medial theta (FMT) power. Cluster-corrected comparison of oscillatory power in the delay period of the 1-back task relative to the same period in the DMS task revealed a significant increase in theta power (4-8 Hz; **A**, summed across significant channels) at fronto-medial channels (**B**, summed across significant frequencies from 4-8 Hz – note the same topography of significant channels emerged when summing across the full 4-9 Hz range).

### Decoding stimulus category during WM maintenance

We next addressed the question of when and where WM content is maintained during the delay period. To this end, multivariate pattern classification was applied to the delay period of object and scene trials of the 1-back task. Specifically, the ability to decode stimulus content was assessed via linear discriminant analysis (LDA) in a k-fold cross-validation regimen using – at each time point – the raw EEG signal across channels as features (see method section for additional details). Classifier accuracy was compared to chance (50%) and corrected across time by cluster-based permutation. As shown in Figure 3A, this comparison revealed an extended interval of significant above-chance decoding during the delay period, i.e., when no stimulus was visually present. Of note, decoding performance was also significantly above chance when averaging across the entire delay period (t_(27)_ = 1.82, p = 0.04). Classification was also conducted on the stimulus and response periods, both of which showed accurate decoding across time (Figure 3A) as well as when collapsing across time: stimulus t_(27)_ = 11.99, p < 0.001; response t_(27)_ = 5.85, p < 0.001).

**Figure 3.**
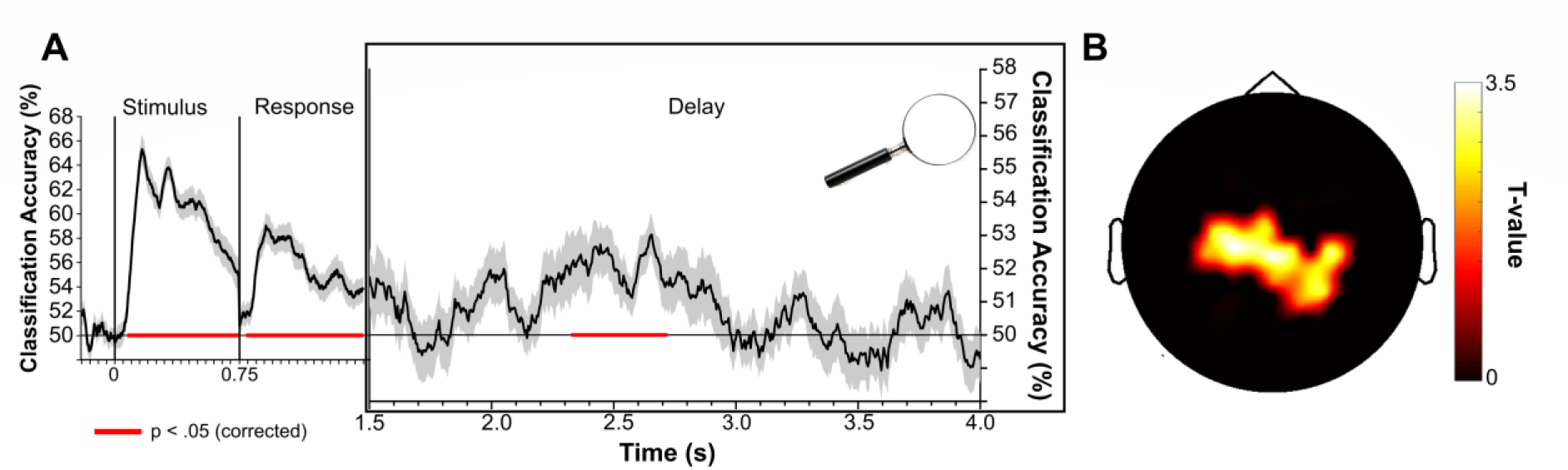
Multivariate decoding of WM content during the 1-back task. **A.** Decoding (mean +/− SEM across participants) of object vs. scene stimuli across time showed significant above-chance (50%) accuracy in all three trial periods (stimulus, response and delay). Depicted trial periods reflect the minimum length, omitting variable jitter durations.
**B.** Spatial searchlight decoding during the significant delay period, revealing maximum performance at parieto-occipital channels (1-tailed t test vs. chance).

Previous findings suggest that WM content is maintained by posterior sensory rather than frontal executive regions (Sreenivasan et al., 2014). We thus repeated the classification analysis with a searchlight approach, specifically during the delay period where no visual information was on-screen. Classification performance was assessed for each channel, including its immediate neighbours (mean number of neighbours = 5.7) and focussing on the period that showed maximal decoding across time when including all channels (880-1305 ms into the delay period; Figure 3A). This approach revealed that stimulus decoding during the delay period was driven largely by parieto-occipital channels (Figure 3B).

Together, these results suggest that stimulus content maintained in WM can be decoded successfully during the delay period of a 1-back task. The parieto-occipital topography of maximal decodability (despite WM-load-related theta changes over fronto-medial channels; Figure 2B) is consistent with findings from fMRI (Harrison & Tong, 2009) and with the notion of frontal theta as an executive control system that does not directly maintain WM content (D’Esposito & Postle, 2015).

### Parietal cortex is functionally coupled with frontal theta

We next probed whether regions involved in the maintenance of WM content (Figure 3) are coupled to frontal theta rhythms (Figure 2). If theta activity does indeed serve as the mediator between frontal executive and posterior representational regions, one would expect increased coherence between these two regions as a result of increased WM engagement. Consequently, we calculated coherence values between channel Fz (representing the centre of the frontal theta cluster previously identified, Figure 2B) and every other channel. Comparison of coherence maps for 1-back vs. DMS tasks revealed a significant centro-parietal cluster of increased coherence during the portion of the delay period when content could be significantly decoded (Figure 4). Of note and as illustrated in the inset of Figure 4, the resulting cluster overlapped markedly with results from our searchlight decoding approach (Figure 3B), suggesting that the same region(s) that maintain WM content is/are coupled to the frontal theta rhythm. Examining the frequency profile of this cluster, it showed maximal coherence at ~7 Hz matching that of the frontal power effect (See Figure 2B). Examining coherence across the full delay period (rather than focusing only on when successful decoding was present), a significant cluster of increased coherence in the 1-back relative to the DMS task was also evident. This cluster closely resembled the previous cluster in both topography and frequency, being located centro-parietally and showing maximal coherence at ~7 Hz.

**Figure 4.**
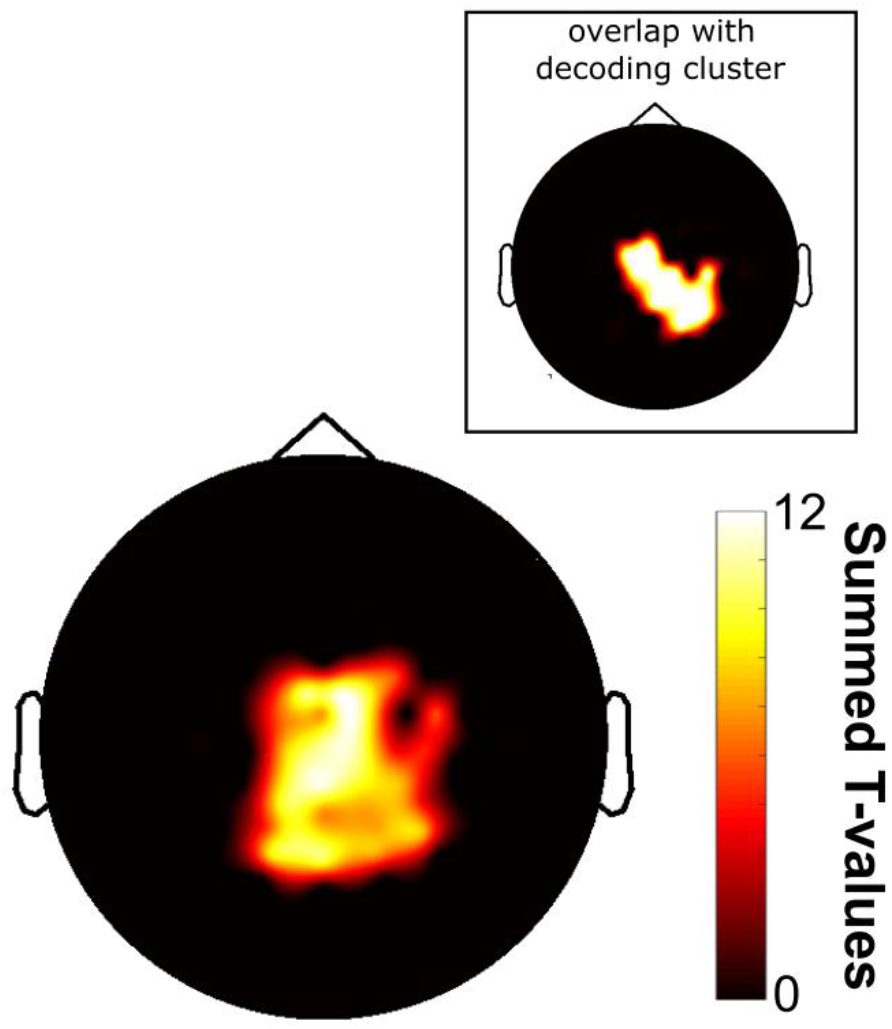
Coupling between posterior ‘content’ channels and frontal theta activity. Taking Fz as the seed channel and comparing 1-back vs. DMS task during the time in the delay period when WM content could be significantly decoded (880-1305 ms) revealed a significant increase in coherence with central-parietal channels in the theta band (4-8 Hz). **Main.** Significant channels, summed across significant frequencies. **Inset.** Overlap with channels contributing to WM content decoding (c.f., Figure 3B).

### Increasing WM load slows down theta

Given the central role of FMT in coordinating WM, how does an increase in WM load impact theta oscillations? One effect of increased WM load might be an increase in FMT power, perhaps reflecting a greater number and/or level of synchronisation of participating neurons (Buzsáki et al., 2012; Buzsáki & Draguhn, 2004). Indeed, such power increases have been reported before (e.g. (Jensen & Tesche, 2002; Meltzer et al., 2008). Another result of increased WM load might be a slowing of the theta rhythm. For instance, the Jensen and Lisman model (Jensen & Lisman, 1998; Lisman & Jensen, 2013) holds that the ongoing theta cycle governs the serial reactivation item-coding cell assemblies. Thus, the duration of a given theta cycle is the limiting factor in how many items can be successfully maintained. A slowing in frequency would therefore facilitate the maintenance additional items within the same theta cycle. In our study, we tested the effect of increased WM load on FMT by comparing the 1-back task with a 2-back variant (Figure 1).

Behaviourally, participants continued to show high accuracy in the 2-back task, although, as with the 1-back task, accuracy was significantly lower in the 2-back relative to the DMS task (t_(27)_ = 3.36, p = 0.002). While there was no significant difference in accuracy between the 2-back and the 1-back tasks (t_(27)_ = 0.816, p = 0.42), the increase in WM load did induce a significant slowing of reaction times (RTs) (paired-samples t-test of 2-back vs. 1-back; t_(27)_ = 3.85, p < 0.001).

We first examined the effects of load (1-back vs. 2-back) and frequency (between 4-8 Hz in 0.5 Hz increments) on power within the theta band over the frontal-medial cluster. Despite previous reports of theta power scaling with WM load (Jensen & Tesche, 2002; Meltzer et al., 2008), we did not find strong evidence for a theta power increase from the 1-back to the 2-back task in the present data. Examining fronto-medial theta power (averaged across the delay period) via a repeated-measures ANOVA revealed no significant main effect of load (F_(1,27)_ = 3.26, p = 0.082). Unsurprisingly, given the 1/f component present in EEG data (Donoghue et al., 2020), there was a main effect of frequency (F_(8,27)_ = 7.56, p < 0.001). Importantly however, there was also a significant interaction between frequency and load (F_(8,216)_ = 9.807, p < 0.001). Follow-up paired t-tests demonstrated that this was driven by a relative power increase in the 2-back condition in the lower theta range (at frequencies between 4 and 6 Hz, [4 Hz, t_(27)_ = 3.29, p = 0.003; 4.5 Hz, t_(27)_ = 3.53, p = 0.002; 5 Hz, t_(27)_ = 3.07, p = 0.005; 5.5 Hz, t_(27)_ = 2.72, p = 0.011; 6 Hz, t_(27)_ = 2.41, p = 0.023]).

To confirm that this change in frequency was indeed a slowing (and thus broadening) of theta oscillations, we defined, for each participant and n-back condition, the peak theta frequency (4-8 Hz) during the delay period. For every trial and for every participant, the frequency at which the most prominent peak in the spectrum occurred was taken. These peak values were averaged by condition resulting in an average theta peak for each condition for each participant. Consistent with the Jensen and Lisman model, a paired t-test revealed a small but highly consistent decrease in peak frequency between the 1-back and 2-back tasks (means: 5.85 Hz vs. 5.77 Hz; t_(27)_ = 5.02, p < 0.001; see Figure 5). This peak detection approach should be largely insensitive to the 1/f component of EEG signals, but to ensure that this was the case, and given the possible functional significance of a change in this exponent (Donoghue et al., 2020), the same method was applied to data to which the IRASA algorithm had been applied (Wen & Liu, 2016). The significant slowing effect persisted [t_(27)_ = 2.84, p = 0.008]. Finally, to rule out the possibility that this finding had occurred due to any volatility in calculating power on the single trial level, the individual trial spectra were smoothed by sliding average (variable window; 0.20-1.00 Hz). Again, significant slowing for 2-back vs. 1-back was observed for all smoothing ranges [t-values ≥ 4.24, p-values < 0.001].

**Figure 5.**
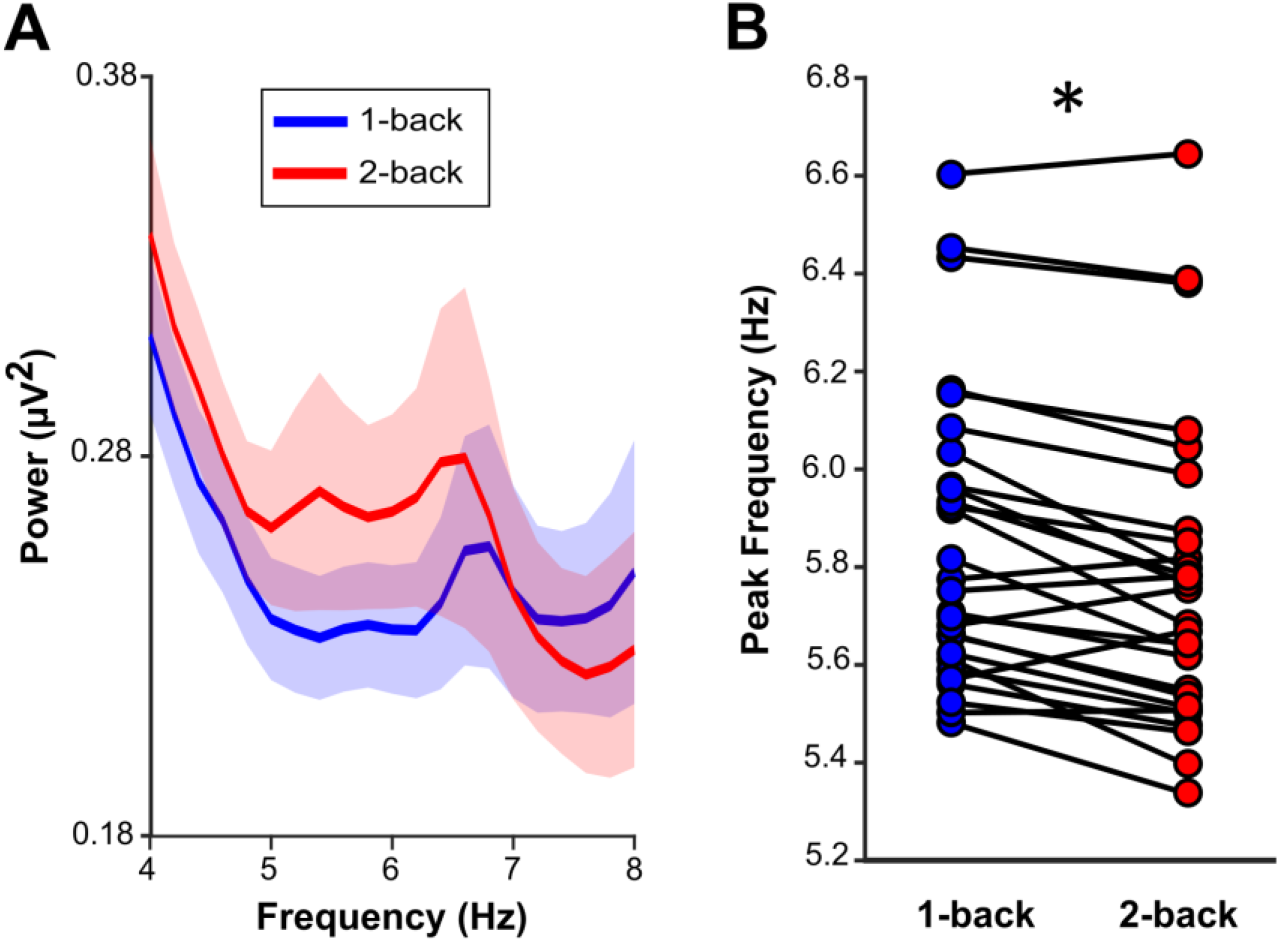
Theta frequency slows to accommodate an additional WM item. Raw Mean (+/− SEM) power values in the theta band (4-8 Hz) across participants, averaged across the delay periods of the 1-back task (blue) and the 2-back task (red). Note the relative shift towards lower frequencies for the 2-back task. **B.** Theta peak frequencies for 1-back and 2-back delay periods shown for each participant. Asterisk indicates statistical significance at α = 0.05.

## Discussion

Our study elucidates the dynamic interplay between the two key components of working memory (WM), i.e., an executive control mechanism and the representation of stimulus content. Using a paradigm that manipulated WM load (Figure 1), we found a power increase in fronto-medial theta (FMT) during 1-back tasks relative to a delayed-match-to-sample (DMS) task (Figure 2). Multivariate pattern analysis showed that WM content, i.e., whether an individual maintained an object or a scene image, could be decoded successfully during the delay period from parieto-occipital channels (rather than from those channels showing the theta power effects, Figure 3). Importantly, channels that contributed to decoding also showed increased coherence in the theta frequency band with frontal sites sensitive to WM load. Lastly, we found that maintaining an additional item in WM leads to slowing of the FMT rhythm (Figure 5), consistent with computational accounts suggesting a broadening of the theta cycle to accommodate multiple WM items.

The finding that WM-induced FMT also governed functional coupling with posterior channels that are most informative to decoding strongly points to a role of FMT in coordinating posterior WM maintenance. This observation unifies a series of recent findings and highlights the importance of theta oscillations as the interlocutor between regional WM sub-systems. Although WM content has been shown to be preferentially decoded from posterior regions, there has been a paucity of evidence to directly associate a measure of WM content with theta activity from frontal regions. That is, there is considerable evidence of FMT as an executive control system in WM (Hsieh & Ranganath, 2014; Riddle et al., 2020) and of posterior localisation of WM content (Christophe et al., 2012; Harrison & Tong, 2009). However, there is little extant evidence directly connecting WM content with frontal theta activity. Previous work has largely focussed on how frontal theta interacts either with activity in other frequency bands, e.g., gamma activity (Berger et al., 2019), or with neuronal spiking (Crowe et al., 2013; Siegel et al., 2009), neither of which provide a direct read-out of high-level WM content. A recent study which did measure WM content observed modulation of decoding from posterior regions according to a theta/alpha rhythm (ten Oever et al., 2020), but did not link this pattern to frontal activity.

The topographical dissociation of frontal control mechanisms (Figure 2B) vs. posterior content maintenance (Figure 3B) dovetails with WM models proposing a domain-general role for prefrontal cortex in executive control (D’Esposito & Postle, 2015). According to the sensory recruitment model, the direct maintenance of content is then accomplished by posterior regions (Scimeca et al., 2018), specifically those that are involved in the processing of the stimulus in a non-WM context. However, the classification we conducted here was between object and scene stimuli. Our data thus do not preclude the possibility that some degree of information is maintained in frontal regions – only that this information does not distinguish between objects vs. scenes as used here. It has, however, been suggested that decoding from frontal regions reflects a transformed stimulus representation in anticipation of a stimulus comparison (Christophel et al., 2017; Lee et al., 2013). In the present case, the stimulus category was orthogonal to whether the stimulus was a match or not (and hence which response to give), ruling out that WM decoding could reflect any planning-related or future response-oriented activity.

Could decodability of WM content during the delay period reflect a spill-over from the preceding stimulus or response interval? In a recent EEG study in which grating orientations were decoded, there was an initial increase in accuracy after stimulus offset followed by a sustained decline (Bae & Luck, 2018). This pattern more closely resembles the response period in our dataset (Figure 3) and is consistent with the finding that decodability rebounds after stimulus offset (Robinson et al., 2019). However, stimulus presentation and maintenance were separated by a minimum of 750 ms in our paradigm (Figure 1) and maximum decodability was actually seen from 880 ms to 1305 ms after delay onset, mitigating impacts of preceding stimulus or response windows.

Assuming that a frontal executive does coordinate WM maintenance via theta oscillations, how does this system respond to increasing task demand? One possibility is a scaling of theta power. Theta power is frequently greater in conditions in which more items must be stored in WM (Gevins et al., 1997; Hsieh & Ranganath, 2014). Furthermore, parametric theta power scaling with conditions of increasing load has previously been observed (Jensen & Tesche, 2002; Meltzer et al., 2008), although not without exception. Payne and Kounios (2009) systematically varied load by presenting 2, 4, or 6 letters, observing an increase in fronto-parietal coherence but not in theta power. Many of these studies employed Sternberg-like paradigms, but studies more comparable to the current paradigm also show some inconsistencies. Brookes et al. (2011) observed robust theta power increases between 0-, 1-, and 2-back tasks, whereas Missonnier et al. (2006) did not find a significant difference between 1- and 2-back conditions. The extent to which the effect of load is influenced by other differences between paradigms, such as stimulus complexity, block length or delay duration, are important to address in future work. In the present study, although FMT power was greater in the 1-back relative to the DMS task, this increase did not extend to the 2-back task.

An alternative way FMT might respond to WM load is a change in frequency. Indeed, in place of a scaling of FMT power, we provide here the first empirical evidence of a slowing of the FMT rhythm in response to increasing WM load. The magnitude of slowing was moderate, but highly consistent across participants. According to the Jensen-Lisman model (Jensen & Lisman, 1998; Lisman & Jensen, 2013), slowing of the carrier theta frequency facilitates bursting of additional item-coding cell assemblies in each cycle whilst maintaining phase separation among items. Although a specific slowing in FMT has not previously been demonstrated, Axmacher et al. (2010) did observe, in intracranial hippocampal recordings during a WM (Sternberg) task, a load-dependent reduction of the theta frequency modulating power in the beta/low gamma band. Further indirect evidence for load-dependent theta slowing comes from a series of studies employing transcranial alternating current stimulation (tACS). Modulating the speed of endogenous theta by means of stimulating at a low (3 Hz) or high (7 Hz) theta frequency was shown to improve or impede WM function, respectively (Bender et al., 2019; Vosskuhl et al., 2015; Wolinski et al., 2018). The data here are in agreement with the implication of these stimulation studies - while theta power is critical for WM (as evidenced by the increase in the 1-back relative to the DMS task), the limiting factor in holding multiple items may in fact be the frequency of ongoing theta oscillations.

It should be noted that these stimulation studies and the present study are agnostic regarding certain specifics of theta phase-coding in WM. Neither differentiates between the theta-gamma framework as originally proposed and a more recent account in which individual WM items are serially reactivated on the peak of distinct theta cycles (Herman et al., 2013; Sauseng et al., 2019). In the latter case, a slower theta oscillation allows for a longer period of time in which a single item can be more robustly reactivated, thereby improving the fidelity of each representation.

To summarise, we show that frontal theta rhythms orchestrate the maintenance of stimulus representations in posterior brain regions in the service of working memory performance. Increasing the amount of information to be maintained led to a slowing of theta frequency, consistent with the idea that longer duty cycles are needed to accommodate additional items held in working memory.

## Acknowledgements

This work was supported by a Wellcome Trust/Royal Society Sir Henry Dale Fellowship (107672/Z/15/Z) to B.P.S. and an MRC IMPACT PhD studentship awarded to O.R. (1973157).

## References

Axmacher, N., Henseler, M. M., Jensen, O., Weinreich, I., Elger, C. E., & Fell, J. (2010). Cross-frequency coupling supports multi-item working memory in the human hippocampus. Proceedings of the National Academy of Sciences of the United States of America. https://doi.org/10.1073/pnas.0911531107

Baddeley, A. (1992). Working memory. Science. https://doi.org/10.1126/science.1736359

Baddeley, A. (2003). Working memory: Looking back and looking forward. Nature Reviews Neuroscience. https://doi.org/10.1038/nrn1201

Bae, G. Y., & Luck, S. J. (2018). Dissociable decoding of spatial attention and working memory from EEG oscillations and sustained potentials. Journal of Neuroscience. https://doi.org/10.1523/JNEUROSCI.2860-17.2017

Bahramisharif, A., Jensen, O., Jacobs, J., & Lisman, J. (2018). Serial representation of items during working memory maintenance at letter-selective cortical sites. PLoS Biology. https://doi.org/10.1371/journal.pbio.2003805

Bastos, A. M., & Schoffelen, J. M. (2016). A tutorial review of functional connectivity analysis methods and their interpretational pitfalls. In Frontiers in Systems Neuroscience. https://doi.org/10.3389/fnsys.2015.00175

Bender, M., Romei, V., & Sauseng, P. (2019). Slow Theta tACS of the Right Parietal Cortex Enhances Contralateral Visual Working Memory Capacity. Brain Topography. https://doi.org/10.1007/s10548-019-00702-2

Berger, B., Griesmayr, B., Minarik, T., Biel, A. L., Pinal, D., Sterr, A., & Sauseng, P. (2019). Dynamic regulation of interregional cortical communication by slow brain oscillations during working memory. Nature Communications. https://doi.org/10.1038/s41467-019-12057-0

Brainard, D. H. (1997). The Psychophysics Toolbox. Spatial Vision. https://doi.org/10.1163/156856897X00357

Bressler, S. L., & Menon, V. (2010). Large-scale brain networks in cognition: emerging methods and principles. In Trends in Cognitive Sciences. https://doi.org/10.1016/j.tics.2010.04.004

Brodeur, M. B., Dionne-Dostie, E., Montreuil, T., & Lepage, M. (2010). The bank of standardized stimuli (BOSS), a new set of 480 normative photos of objects to be used as visual stimuli in cognitive research. PLoS ONE. https://doi.org/10.1371/journal.pone.0010773

Brookes, M. J., Wood, J. R., Stevenson, C. M., Zumer, J. M., White, T. P., Liddle, P. F., & Morris, P. G. (2011). Changes in brain network activity during working memory tasks: A magnetoencephalography study. NeuroImage. https://doi.org/10.1016/j.neuroimage.2010.10.074

Buzsáki, G., Anastassiou, C. A., & Koch, C. (2012). The origin of extracellular fields and currents-EEG, ECoG, LFP and spikes. In Nature Reviews Neuroscience. https://doi.org/10.1038/nrn3241

Buzsáki, G., & Draguhn, A. (2004). Neuronal olscillations in cortical networks. In Science. https://doi.org/10.1126/science.1099745

Christophe, T. B., Hebart, M. N., & Haynes, J. D. (2012). Decoding the contents of visual short-term memory from human visual and parietal cortex. Journal of Neuroscience. https://doi.org/10.1523/JNEUROSCI.0184-12.2012

Christophel, T. B., Klink, P. C., Spitzer, B., Roelfsema, P. R., & Haynes, J. D. (2017). The Distributed Nature of Working Memory. In Trends in Cognitive Sciences. https://doi.org/10.1016/j.tics.2016.12.007

Crowe, D. A., Goodwin, S. J., Blackman, R. K., Sakellaridi, S., Sponheim, S. R., MacDonald, A. W., & Chafee, M. V. (2013). Prefrontal neurons transmit signals to parietal neurons that reflect executive control of cognition. Nature Neuroscience. https://doi.org/10.1038/nn.3509

D’Esposito, M., & Postle, B. R. (2015). The Cognitive Neuroscience of Working Memory. Annual Review of Psychology. https://doi.org/10.1146/annurev-psych-010814-015031

Donoghue, T., Haller, M., Peterson, E. J., Varma, P., Sebastian, P., Gao, R., Noto, T., Lara, A. H., Wallis, J. D., Knight, R. T., Shestyuk, A., & Voytek, B. (2020). Parameterizing neural power spectra into periodic and aperiodic components. Nature Neuroscience. https://doi.org/10.1038/s41593-020-00744-x

Gevins, A., Smith, M. E., McEvoy, L., & Yu, D. (1997). High-resolution EEG mapping of cortical activation related to working memory: Effects of task difficulty, type of processing, and practice. Cerebral Cortex. https://doi.org/10.1093/cercor/7.4.374

Harrison, S. A., & Tong, F. (2009). Decoding reveals the contents of visual working memory in early visual areas. Nature. https://doi.org/10.1038/nature07832

Herman, P. A., Lundqvist, M., & Lansner, A. (2013). Nested theta to gamma oscillations and precise spatiotemporal firing during memory retrieval in a simulated attractor network. Brain Research. https://doi.org/10.1016/j.brainres.2013.08.002

Hsieh, L. T., & Ranganath, C. (2014). Frontal midline theta oscillations during working memory maintenance and episodic encoding and retrieval. In NeuroImage. https://doi.org/10.1016/j.neuroimage.2013.08.003

Imaruoka, T., Saiki, J., & Miyauchi, S. (2005). Maintaining coherence of dynamic objects requires coordination of neural systems extended from anterior frontal to posterior parietal brain cortices. NeuroImage. https://doi.org/10.1016/j.neuroimage.2005.01.045

Jensen, O., & Lisman, J. E. (1998). An oscillatory short-term memory buffer model can account for data on the Sternberg task. Journal of Neuroscience. https://doi.org/10.1523/jneurosci.18-24-10688.1998

Jensen, O., & Tesche, C. D. (2002). Frontal theta activity in humans increases with memory load in a working memory task. European Journal of Neuroscience. https://doi.org/10.1046/j.1460-9568.2002.01975.x

Kearney, K. (2019). boundedline (https://uk.mathworks.com/matlabcentral/fileexchange/27485-boundedline-m). Retrieved 2019.

Kleiner, M., Brainard, D. H., Pelli, D. G., Broussard, C., Wolf, T., & Niehorster, D. (2007). What’s new in Psychtoolbox-3? Perception. https://doi.org/10.1068/v070821

Kuczenski, B. (2019). hline and vline (https://www.mathworks.com/matlabcentral/fileexchange/1039-hline-and-vline), MATLAB Central File Exchange. Retrieved 2019.

Lee, S. H., Kravitz, D. J., & Baker, C. I. (2013). Goal-dependent dissociation of visual and prefrontal cortices during working memory. Nature Neuroscience. https://doi.org/10.1038/nn.3452

Lisman, J. E., & Jensen, O. (2013). The Theta-Gamma Neural Code. In Neuron. https://doi.org/10.1016/j.neuron.2013.03.007

Maris, E., & Oostenveld, R. (2007). Nonparametric statistical testing of EEG- and MEG-data. Journal of Neuroscience Methods. https://doi.org/10.1016/j.jneumeth.2007.03.024

Maurer, U., Brem, S., Liechti, M., Maurizio, S., Michels, L., & Brandeis, D. (2014). Frontal Midline Theta Reflects Individual Task Performance in a Working Memory Task. Brain Topography. https://doi.org/10.1007/s10548-014-0361-y

Meltzer, J. A., Zaveri, H. P., Goncharova, I. I., Distasio, M. M., Papademetris, X., Spencer, S. S., Spencer, D. D., & Constable, R. T. (2008). Effects of working memory load on oscillatory power in human intracranial EEG. Cerebral Cortex. https://doi.org/10.1093/cercor/bhm213

Michels, L., Bucher, K., Lüchinger, R., Klaver, P., Martin, E., Jeanmonod, D., & Brandeis, D. (2010). Simultaneous EEG-fMRI during a working memory task: Modulations in low and high frequency bands. PLoS ONE. https://doi.org/10.1371/journal.pone.0010298

Missonnier, P., Deiber, M. P., Gold, G., Millet, P., Gex-Fabry Pun, M., Fazio-Costa, L., Giannakopoulos, P., & Ibáñez, V. (2006). Frontal theta event-related synchronization: Comparison of directed attention and working memory load effects. Journal of Neural Transmission. https://doi.org/10.1007/s00702-005-0443-9

Onton, J., Delorme, A., & Makeig, S. (2005). Frontal midline EEG dynamics during working memory. NeuroImage. https://doi.org/10.1016/j.neuroimage.2005.04.014

Oostenveld, R., Fries, P., Maris, E., & Schoffelen, J. M. (2011). FieldTrip: Open source software for advanced analysis of MEG, EEG, and invasive electrophysiological data. Computational Intelligence and Neuroscience. https://doi.org/10.1155/2011/156869

Owen, A. M., McMillan, K. M., Laird, A. R., & Bullmore, E. (2005). N-back working memory paradigm: A meta-analysis of normative functional neuroimaging studies. Human Brain Mapping. https://doi.org/10.1002/hbm.20131

Payne, L., & Kounios, J. (2009). Coherent oscillatory networks supporting short-term memory retention. Brain Research. https://doi.org/10.1016/j.brainres.2008.09.095

Polanía, R., Nitsche, M. A., Korman, C., Batsikadze, G., & Paulus, W. (2012). The importance of timing in segregated theta phase-coupling for cognitive performance. Current Biology. https://doi.org/10.1016/j.cub.2012.05.021

R Core Team. (2018). A Language and Environment for Statistical Computing. In R Foundation for Statistical Computing.

Raghavachari, S., Kahana, M. J., Rizzuto, D. S., Caplan, J. B., Kirschen, M. P., Bourgeois, B., Madsen, J. R., & Lisman, J. E. (2001). Gating of human theta oscillations by a working memory task. Journal of Neuroscience. https://doi.org/10.1523/JNEUROSCI.21-09-03175.2001

Reinhart, R. M. G., & Nguyen, J. A. (2019). Working memory revived in older adults by synchronizing rhythmic brain circuits. Nature Neuroscience. https://doi.org/10.1038/s41593-019-0371-x

Riddle, J., Scimeca, J. M., Cellier, D., Dhanani, S., & D’Esposito, M. (2020). Causal Evidence for a Role of Theta and Alpha Oscillations in the Control of Working Memory. Current Biology. https://doi.org/10.1016/j.cub.2020.02.065

Robinson, A. K., Grootswagers, T., & Carlson, T. A. (2019). The influence of image masking on object representations during rapid serial visual presentation. NeuroImage. https://doi.org/10.1016/j.neuroimage.2019.04.050

Sauseng, P., Griesmayr, B., Freunberger, R., & Klimesch, W. (2010). Control mechanisms in working memory: A possible function of EEG theta oscillations. In Neuroscience and Biobehavioral Reviews. https://doi.org/10.1016/j.neubiorev.2009.12.006

Sauseng, P., Klimesch, W., Doppelmayr, M., Hanslmayr, S., Schabus, M., & Gruber, W. R. (2004). Theta coupling in the human electroencephalogram during a working memory task. Neuroscience Letters. https://doi.org/10.1016/j.neulet.2003.10.002

Sauseng, P., Peylo, C., Biel, A. L., Friedrich, E. V. C., & Romberg-Taylor, C. (2019). Does cross-frequency phase coupling of oscillatory brain activity contribute to a better understanding of visual working memory? British Journal of Psychology. https://doi.org/10.1111/bjop.12340

Scimeca, J. M., Kiyonaga, A., & D’Esposito, M. (2018). Reaffirming the Sensory Recruitment Account of Working Memory. In Trends in Cognitive Sciences. https://doi.org/10.1016/j.tics.2017.12.007

Siegel, M., Warden, M. R., & Miller, E. K. (2009). Phase-dependent neuronal coding of objects in short-term memory. Proceedings of the National Academy of Sciences of the United States of America. https://doi.org/10.1073/pnas.0908193106

Sreenivasan, K. K., Curtis, C. E., & D’Esposito, M. (2014). Revisiting the role of persistent neural activity during working memory. In Trends in Cognitive Sciences. https://doi.org/10.1016/j.tics.2013.12.001

ten Oever, S., De Weerd, P., & Sack, A. T. (2020). Phase-dependent amplification of working memory content and performance. Nature Communications. https://doi.org/10.1038/s41467-020-15629-7

Treder, M. S. (2020). MVPA-Light: A Classification and Regression Toolbox for Multi-Dimensional Data. Frontiers in Neuroscience. https://doi.org/10.3389/fnins.2020.00289

Tsujimoto, T., Shimazu, H., & Isomura, Y. (2006). Direct recording of theta oscillations in primate prefrontal and anterior cingulate cortices. Journal of Neurophysiology. https://doi.org/10.1152/jn.00730.2005

Vogel, E. K., & Machizawa, M. G. (2004). Neural activity predicts individual differences in visual working memory capacity. Nature. https://doi.org/10.1038/nature02447

Von Stein, A., & Sarnthein, J. (2000). Different frequencies for different scales of cortical integration: From local gamma to long range alpha/theta synchronization. International Journal of Psychophysiology. https://doi.org/10.1016/S0167-8760(00)00172-0

Vosskuhl, J., Huster, R. J., & Herrmann, C. S. (2015). Increase in short-term memory capacity induced by down-regulating individual theta frequency via transcranial alternating current stimulation. Frontiers in Human Neuroscience. https://doi.org/10.3389/fnhum.2015.00257

Wen, H., & Liu, Z. (2016). Separating Fractal and Oscillatory Components in the Power Spectrum of Neurophysiological Signal. Brain Topography. https://doi.org/10.1007/s10548-015-0448-0

Wolinski, N., Cooper, N. R., Sauseng, P., & Romei, V. (2018). The speed of parietal theta frequency drives visuospatial working memory capacity. PLoS Biology. https://doi.org/10.1371/journal.pbio.2005348

Xiao, J., Hays, J., Ehinger, K. A., Oliva, A., & Torralba, A. (2010). SUN database: Large-scale scene recognition from abbey to zoo. Proceedings of the IEEE Computer Society Conference on Computer Vision and Pattern Recognition. https://doi.org/10.1109/CVPR.2010.5539970

Zakrzewska, M. Z., & Brzezicka, A. (2014). Working memory capacity as a moderator of load-related frontal midline theta variability in Sternberg task. Frontiers in Human Neuroscience. https://doi.org/10.3389/fnhum.2014.00399

Zuure, M. B., Hinkley, L. B. N., Tiesinga, P. H. E., Nagarajan, S. S., & Cohen, M. X. (2020). Multiple midfrontal thetas revealed by source separation of simultaneous MEG and EEG. BioRxiv. https://doi.org/10.1101/2020.03.11.987040

